# Gaze Behaviour During Sensorimotor Adaptation Parcellates the Explicit and Implicit Contributions to Learning

**DOI:** 10.1101/237651

**Authors:** Anouk J. de Brouwer, Mohammed Albaghdadi, J. Randall Flanagan, Jason P. Gallivan

## Abstract

Successful motor performance relies on our ability to adapt to changes in the environment by learning novel mappings between motor commands and sensory outcomes. Such adaptation is thought to involve two distinct mechanisms: An implicit, error-based component linked to slow learning and an explicit, strategic component linked to fast learning and savings (i.e., faster relearning). Because behaviour, at any given moment, is the resultant combination of these two processes, it has remained a challenge to parcellate their relative contributions to performance. The explicit component to visuomotor rotation (VMR) learning has recently been measured by having participants verbally report their aiming strategy used to counteract the rotation. However, this procedure has been shown to magnify the explicit component. Here we tested whether task-specific eye movements, a natural component of reach planning—but poorly studied in motor learning tasks—can provide a direct read-out of the state of the explicit component during VMR learning. We show, by placing targets on a visible ring and including a delay between target presentation and reach onset, that individual differences in gaze patterns during sensorimotor adaptation are linked to participants’ rates of learning and can predict the expression of savings. Specifically, we find that participants who, during reach planning, naturally fixate an aimpoint, rotated away from the target location, show faster initial adaptation and readaptation 24 hrs. later. Our results demonstrate that gaze behaviour can not only uniquely identify individuals who implement cognitive strategies during learning, but also how their implementation is linked to differences in learning.

## Introduction

Skilled motor behaviour requires the ability to adapt to changes in the environment that alter the mapping between motor commands and their sensory consequences (Shadmehr et al., 2010; Wolpert et al., 2011). Such adaptation has been extensively investigated using reaching or throwing tasks with displacing prisms (Martin et al., 1996; Bedford, 1999; Fernández-Ruiz and Díaz, 1999; Redding and Wallace, 2006) and reaching tasks under a visuomotor rotation (VMR), in which the viewed position of the hand (or cursor representing the hand) is rotated about the hand start location (e.g., Krakauer et al., 2000, 2005; Wigmore et al., 2002). Traditionally, learning in such tasks was presumed to be driven by an implicit process involving the gradual updating of an internal model, which links motor commands and sensory outcomes, based on errors between predicted and viewed consequences of action (Shadmehr et al., 2010; Wolpert et al., 2011). Several studies, however, have demonstrated that learning can also be augmented by (or interfered with) the use of cognitive strategies (Redding and Wallace, 1993, 2002; Martin et al., 1996; Bock et al., 2003; Mazzoni and Krakauer, 2006; Benson et al., 2011; Fernandez-Ruiz et al., 2011; Taylor and Ivry, 2011). To dissociate the implicit and strategic components of VMR learning, Taylor and colleagues (2014) recently developed a task in which participants, prior to each reaching movement, verbally reported their aiming direction—used to counteract the rotation—via numbers placed on a circle surrounding the hand start position. They demonstrated that learning is the resultant combination of two separate processes: A fast explicit process reflecting strategic aiming, and a more gradual, implicit process reflecting updating of an internal model.

More recently, this verbal reporting task has also been used to probe the mechanisms underlying *savings,* which refers to faster relearning of a previously forgotten (or ‘washed out’) memory (Ebbinghaus, 1913; Brashers-Krug et al., 1996; Krakauer et al., 2005). Morehead et al. (2015) showed that improvements in aiming strategy underlie the faster rate of learning observed when individuals re-encounter the VMR following washout of initial learning. This result, along with the finding that fast learning and relearning are not observed when the expression of the explicit component is mitigated by limiting preparation time (Fernandez-Ruiz et al., 2011; Haith et al., 2015; Leow et al., 2017), suggests that savings are largely driven by the recall of previously implemented strategies. However, because the declarative nature of the verbal reporting task has been shown to influence the explicit (Taylor et al., 2014) or implicit (Taylor et al., 2014; Leow et al., 2017) contributions to learning, alternative measures may be critical to parcelling out their unique contributions to learning and how they shape individual performance.

Eye movements are a fundamental component to the planning and control of visually guided actions (Land and Furneaux, 1997; Johansson et al., 2001). During reach planning, gaze is naturally directed to the target before initiation of the hand movement to improve spatial localization of the target and help guide the hand to the target using visual feedback (e.g., Prablanc et al., 1979; Paillard, 1982). Since the explicit component of VMR adaptation involves strategically re-aiming the hand towards an aimpoint that is rotated away from the target, it is plausible that eye movements are used to identify this aimpoint location. While there is some evidence to suggest that gaze behaviour may be linked to the explicit component of learning (Rand and Rentsch, 2015, 2016), this relationship has not been directly examined, nor has it been explored how the time course of gaze behaviour during learning may be linked to individual differences in learning rates and the expression of savings. Here we tested the novel hypothesis that task-specific eye movements, during a VMR task in which targets are presented on a ring of visual landmarks, can provide a direct ‘readout’ of both the implementation and state of the explicit component over the time course of sensorimotor learning and relearning following washout.

## Methods

### Participants

A total of 56 young right-handed adults participated in one of three experiments. Twenty-one people took part in the Intermittent Report experiment (Experiment 1; 5 men and 16 women, age 18-25 years), after exclusion of two participants due to technical problems. The No Report experiment (Experiment 2) was performed by 21 different participants (8 men and 13 women, age 18-22 years). Twelve participants were recruited for the No Preview experiment (Experiment 3; 5 men and 7 women; age 19-24 years). Participants had normal or corrected-to-normal vision and provided written informed consent before participation. The experiment was part of a research project that was approved by the general research ethics board from Queen’s University.

### Apparatus

Participants were seated at a table and performed center-out reaching movements to visual targets by sliding a stylus across a digitizing tablet (Figure 1A). Stimuli were presented on a vertical LCD monitor (display size 47.5 × 26.5 cm, resolution 1920 × 1080 pixels, refresh rate 60 Hz) placed ~50 cm in front of a chin and forehead rest. Vision of the tablet and hand was occluded by a rectangular piece of black styrofoam attached horizontally below the chin rest. Movement trajectories were sampled at 100 Hz by the digitizing tablet (active area 311 × 216 mm, Wacom Intuous). The ratio between movement of the tip of the stylus and movement of the cursor presented on the screen was set to 1:2, so that a movement of 5 cm on the tablet corresponded to a 10 cm movement of the cursor. Eye movements were tracked at 500 Hz using a video-based eye tracker (Eyelink 1000, SR Research) placed beneath the monitor.

**Figure 1.**
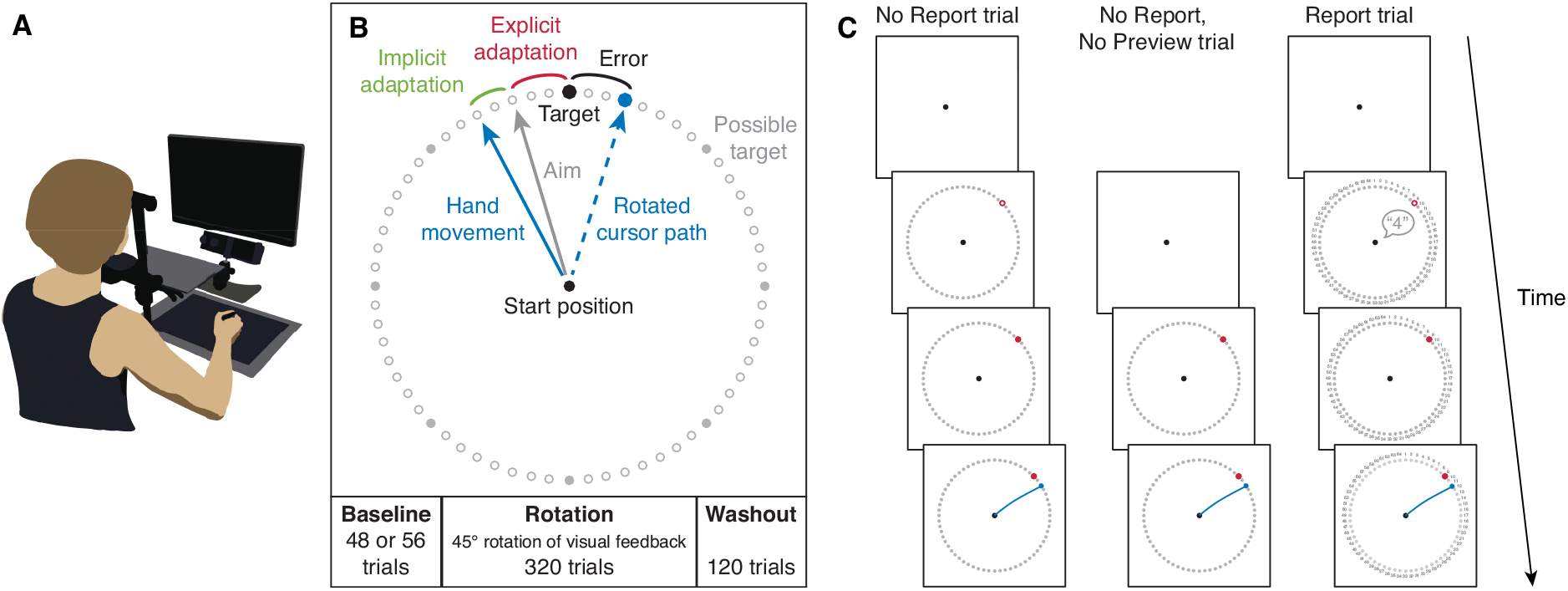
Experimental setup and procedures. **A.** Experimental Setup. Participants performed fast reaching movements by sliding a pen across a digitizing tablet, without vision of the hand. Visual stimuli and the cursor representing the hand position were presented on a monitor. **B.** Task. A target was presented in one of eight locations, and flanked by a ring of landmark circles. Veridical cursor feedback was provided in the baseline and washout blocks. In the rotation block, participants were exposed to a 45° rotation of the cursor feedback. **C.** Trial types. In No Report trials participants were given a 2 s preview of the target and landmarks before the response was cued. In Report trials participants reported their aiming direction via the numbered visual landmarks.

### Procedure

Each trial started with the participant moving the cursor (4 mm radius cyan circle) into the starting position (5 mm radius white circle) using the stylus. The cursor became visible when its center was within 2 cm of the center of the start position. After the cursor was held within the start position for 500 ms, a red target circle (5 mm radius) and 64 outlined grey ‘landmark’ circles (3 mm radius, spaced 5.625° apart) were presented on a ring with a radius of 10 cm (Figure 1B) after a 100 ms delay. The target was presented at one of eight locations, separated by 45° (0, 45, 90, 135, 180, 225, 270 and 315°), in randomized sets of eight trials. As outlined below, the subsequent trial events depended on the trial type.

In no-report trials (used in Experiments 1 and 2), the target initially appeared as an outlined circle, and participants were given a target preview of 2 s before the target filled in, which served as the cue for participants to initiate their reach. In no-report, no-preview trials (used in Experiment 3), the target appeared as a filled circle and participants were instructed to initiate their reach immediately when the target appeared. In report trials (used in Experiment 1), the target was an outlined circle and the visual landmarks were numbered. Participants were required to verbally report the number of the landmark they planned to reach toward for the cursor to hit the target (as in Taylor et al., 2014) the experimenter recorded the number using a keyboard. The target turned red two seconds after its appearance, or immediately after the experimenter recorded the response if the response took longer than two seconds, providing the go signal for the participant to initiate their reach.

In all trials, participants were instructed to make a fast reaching movement, ‘slicing’ through the target. If the movement was initiated (i.e., the cursor had moved fully out of the start circle) before the go cue, the trial was aborted and a feedback text message “Too early” appeared centrally on the screen. In trials with correct timing, the cursor was visible during the movement to the ring (at 10 cm distance) and then became stationary for one second when it reached the ring, providing the participant with visual feedback of their endpoint reach error. If any part of the stationary cursor overlapped with any part of the target, the target was coloured green and the participant received one point. Points were displayed on the screen every 80 trials in the rotation and washout blocks, followed by a 30 s break.

Each testing session took about 75 minutes to complete and consisted of a baseline block with veridical cursor feedback, a rotation block in which feedback of the cursor during the reach was rotated clockwise by 45°, and a ‘washout’ block in which veridical cursor feedback was restored. Participants in Experiments 1 and 2 completed two sessions, separated by a day, whereas participants in Experiment 3 completed a single session. Participants were not informed about the visuomotor rotation.

## Experiment 1: Intermittent Report

In the baseline block, participants first completed 48 no-report trials followed by 8 report trials. In the rotation block, participants completed 320 trials (40 sets of 8 trials). To test whether gaze fixations prior to executing a reach movement can provide a readout of the explicit component of visuomotor adaptation, in the rotation block we randomly intermixed two report trials and six no-report trials in each set of eight trials. This intermittent reporting was introduced after an initial set of eight no-report trials. At this moment, participants were told by the experimenter that “they had probably noticed something strange is going on” and they were instructed to report the direction of their hand movement (not the cursor movement) required to hit the target when the numbers are displayed. In the washout block following the rotation block, participants completed 120 no-report trials without a rotation. To examine savings when re-exposed to the visuomotor rotation, and its relation to gaze patterns, participants performed two identical testing sessions separated by exactly 24 hours.

## Experiment 2: No Report

The second experiment was designed to test the extent to which the implementation of an aiming strategy, and the occurrence of fixations at the aimpoint, is influenced by having participants report their aiming direction. This experiment was identical to the Intermittent Report experiment (Experiment 1) except that the baseline block only included 48 no-report trials, and all 320 trials in the rotation block were no-report trials. To examine savings when re-exposed to the visuomotor rotation, participants performed two identical testing sessions separated by 24 hours.

## Experiment 3: No Preview

We tested a third group of participants to examine the extent to which the implementation of an aiming strategy, and the occurrence of fixations at the aimpoint, depends on having a target preview period, as previous studies have shown strategic aiming is effortful, especially at short preparation times (Leow et al., 2017). The experiment was the same as the No Report experiment (Experiment 2) except that all of the trials in the baseline, rotation, and washout blocks were no-report, no-preview trials, and participants only performed a single testing session. Our instructions did not stress reaction time, but they emphasized that participants had to make a single, fast, uncorrected reaching movement slicing through the target. If the duration of the reach was longer than 400 ms, the trial was aborted and a text feedback message “Too slow” appeared centrally on the screen.

### Data Analysis

#### Hand movements

Trials in which the timing was incorrect (as detected online) were excluded from the offline analysis of hand and eye movements. We also excluded trials in which the movement time, defined as the time between the moment the cursor had fully moved out of the start position until the cursor reached the 10 cm target distance, was longer than 400 ms. To assess task performance on each trial, we calculated the hand angle with respect to the target angle at the moment the cursor reached the target distance. To do this, we first linearly interpolated the position of the cursor to 1000 Hz, then converted its horizontal and vertical position at 10 cm distance from the start position to an angle, and finally subtracted the target angle. The endpoint hand angles were averaged across sets of eight trials, containing one repetition of each target direction.

#### Eye movements

For the Intermittent Report experiment (Experiment 1), we first excluded report trials from the analysis of gaze data, since in these trials participants would naturally direct their eyes to the number they want to report. For all experiments, we excluded trials in which there was missing gaze data during at least 50% of the time from the onset of the target until the cursor crossed the ring (i.e, the preview and movement phases; ~7% of trials in the Intermittent Report experiment; ~8% of trials in the No Report experiment; ~4% of trials in the No Preview experiment). Our analysis focused on participants’ fixation locations during the preview and movement phases. For each trial, we first detected and removed blinks from the x and *y* gaze positions that were provided by the eye tracker. Gaze data were low-pass filtered using a 2^nd^ order recursive Butterworth filter with a cutoff frequency of 50 Hz. The filtered x and *y* gaze positions were used to calculate horizontal, vertical and resultant gaze velocity. To obtain fixations, we first identified saccades as having a resultant velocity of 20 cm/s for 5 or more consecutive samples (10 ms). Saccade onset was defined as the last of 5 samples below the threshold of 20 cm/s, and saccade offset was defined as the first of 5 samples below this threshold. Next, fixations were defined as periods of 50 or more consecutive samples (100 ms) in which a saccade with a minimal displacement of 0.5 cm did not occur. We computed the mean x and *y* gaze positions for each fixation, and converted this to a distance from the start position and an angle relative to the target.

We used the resulting fixation locations to quantify gaze patterns (1) over the time course of a single trial, and (2) over the course of each testing session. To examine gaze patterns over the time course of a trial, we first normalized time by scaling each phase (target preview, reaction time, reach, and feedback) of each trial trial to the mean duration of that phase. Next, we computed, for each participant and each sample of trials in the rotation block, the probability that a fixation occurred in three areas: (1) the start point area (<75% of target distance), (2) the visual target area (75-125% of target distance and within 8.4° of the target angle, thereby including one landmark on each side of the target), and (3) a wide ‘aim area’ between the visual target area and -45°, i.e., the hand angle that would fully counteract the rotation, hereafter called the ‘hand target’. Fixations at locations outside these three areas were very rare.

To examine task-relevant gaze fixations over the course of the testing session, we only used fixation angles between 75 and 125% of the target distance. During the preview period on rotation trials, gaze typically shifted to the visual target briefly after its appearance, and from there gaze shifted, often over two or three saccades, towards the hand target (see Figure 2A and B for an example). Therefore, we selected the fixation angle closest to the hand target, discarding fixations within the target area, to obtain a single measure of the putative ‘aimpoint fixation angle’ for each trial. The darker colored dots in the third column of Figure 3 show the fixations selected using this procedure. For group analyses, the resulting fixation angles were averaged across sets of eight trials, for each set that contained at least two ‘aimpoint fixations’.

**Figure 2.**
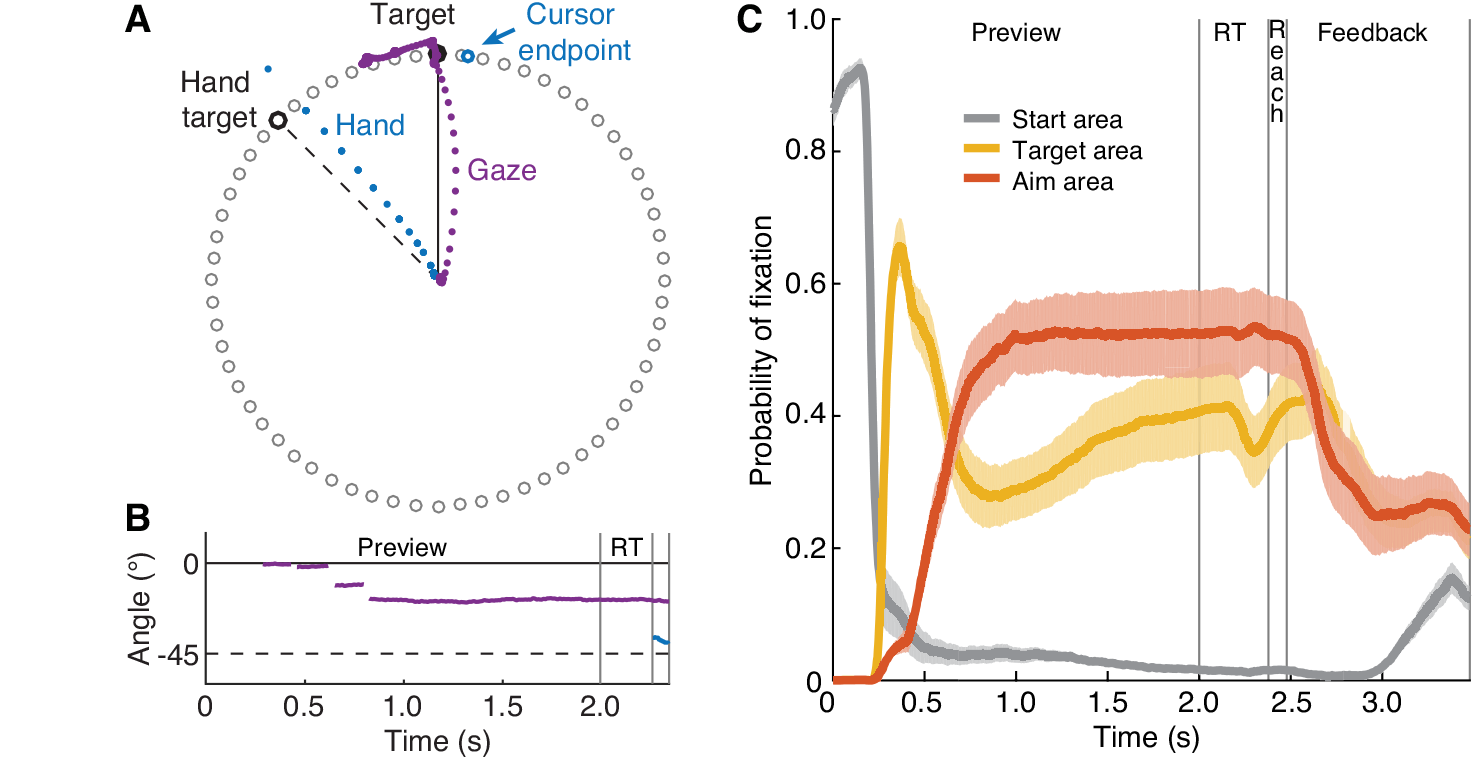
Gaze behaviour in rotation trials. **A**. Typical behaviour in a No Report trial in the rotation block. This participant first moved their gaze to the visual target, and then toward their aiming direction before executing the reach movement. **B**. Time course of fixations (75-125% of target distance; purple), and hand movement (blue) during the target preview, hand reaction time (RT) and reach for the trial shown in A. C. Probability of fixation in the start area (<75% of target distance; gray trace), target area (75-125% of target distance and <8.4° of the visual target; yellow trace), and aim area (75-125% of target distance and -8.4° to -45° from the visual target; orange trace) as a function of within trial timing for the rotation block, averaged across the subgroup of aimpoint fixators in Experiment 1. Shaded areas represent standard error of the mean.

**Figure 3.**
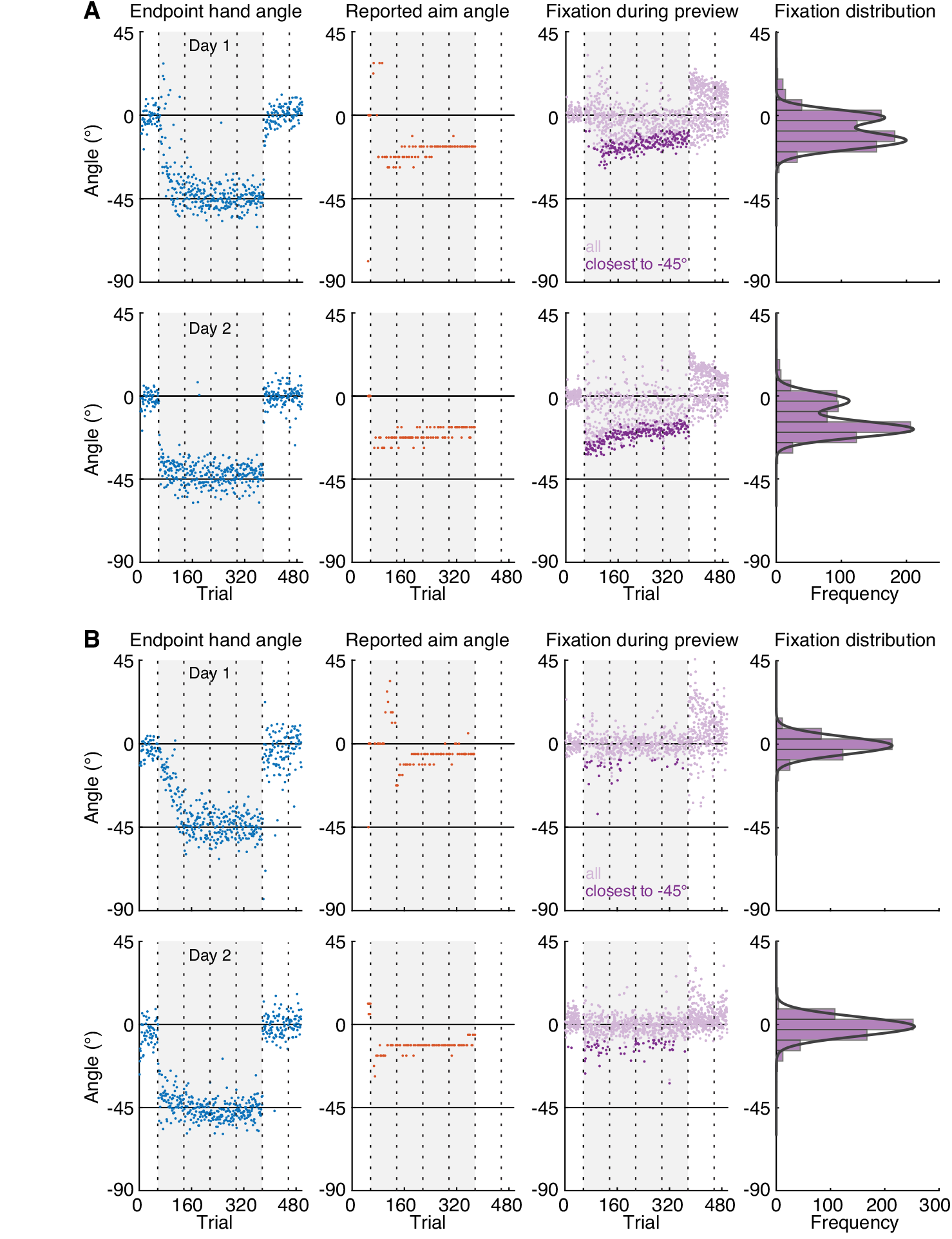
Raw data and experimental approach of classifying participants. **A**. Raw endpoint hand angles (blue), reported aim angles (orange) and fixation angles (purple) during the 2 s target preview period in No Report trials of a representative participant in the Intermittent Report Experiment on day 1 (top) and day 2 (bottom). The grey background indicates when a 45° rotation was applied to the cursor feedback. Vertical dotted lines indicate the timing of 30 s breaks during the experiment. The darker purple dots show, for each trial, the selected fixation angle closest to the hand target, used to compute the group average ‘aimpoint fixation’ angle. The rightmost column shows a histogram of the fixation angles in the rotation block. This participant was classified as an ‘aimpoint fixator’ because their gaze distribution was well fit by a mixture of two Gaussian curves. **B**. Raw data from a representative ‘target-only fixator’, plotted and computed the same as in A. This participant was classified as such because their distribution of fixation angles was unimodal (i.e., fitting procedure yielded a single Gaussian).

#### Gaussian model fitting

Our hypothesis that gaze patterns can provide a readout of the explicit component predicts that the distribution of each participant’s fixation locations should be bimodal, with a peak at the angle of the visual target, and a second peak at the participant’s putative aiming angle. A peak at the aiming angle occurred in the majority, but not all participants. To test for possible differences in learning curves between participants that did or did not exhibit aimpoint fixations, we divided our participants into subgroups of ‘aimpoint fixators’ and ‘target-only fixators’. To do this, we first created, for each participant in Experiment 1 and 2, a histogram of all fixation angles at 75 to 125% target distance during the preview phase of the trials in the rotation block (see Figure 3), excluding the first 40 trials wherein the explicit component changes rapidly (see Figure 4; Taylor et al., 2014). The center of the histogram bins corresponded to the angles of the landmarks, and the width of the bins corresponded to the angular distance in between each two landmarks, such that each bin was 5.625° wide. We used the ‘fit’ function in the MATLAB curve fitting toolbox to perform a nonlinear least squares fit of a mixture of two Gaussian curves to the bin counts y, according to:

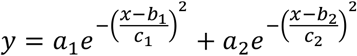

where *a*_1_ and *a*_2_ are the amplitudes of the Gaussians, *b*_1_ and *b*_2_ are the means of the Gaussians and *c*_1_ and *c*_2_ are related to the width of the Gaussians. The lower bounds of the *a*, *b* and *c* parameters were set to [0 -180 0], and the upper bounds were set to [Inf 180 Inf]. The starting value for *a* was set to half of the total bin count, and the starting value for *c* was set to 6 based on initial, unconstrained fits. We set the starting value for *b*_1_ to zero (i.e., the visual target), and for *b*_2_ we used starting values around the mean of the reported aiming direction in the Intermittent Report task (mean±SD: -23.3±7.6; starting values [-30, -28, -26, - 24, -22, -20, -18, -16]). We selected the fit with the highest variance explained by the model. Participants were categorized as ‘aimpoint-fixators’ if the fitting procedure returned two significant Gaussians (see Figure 3A); that is, the 95% of the confidence interval of the means b1 and b2 did not overlap, and the confidence interval of *b*_2_ was outside of the center histogram bin. Otherwise, participants were categorized as target-only fixators, in which case a single Gaussian curve was fit to the bin counts (see Figure 3B). For three participants in Experiments 1 and 2 the confidence interval of the mean of the best fit unimodal curve for one of the days was outside of the center bin. These participants were categorized as aimpoint fixators.

**Figure 4.**
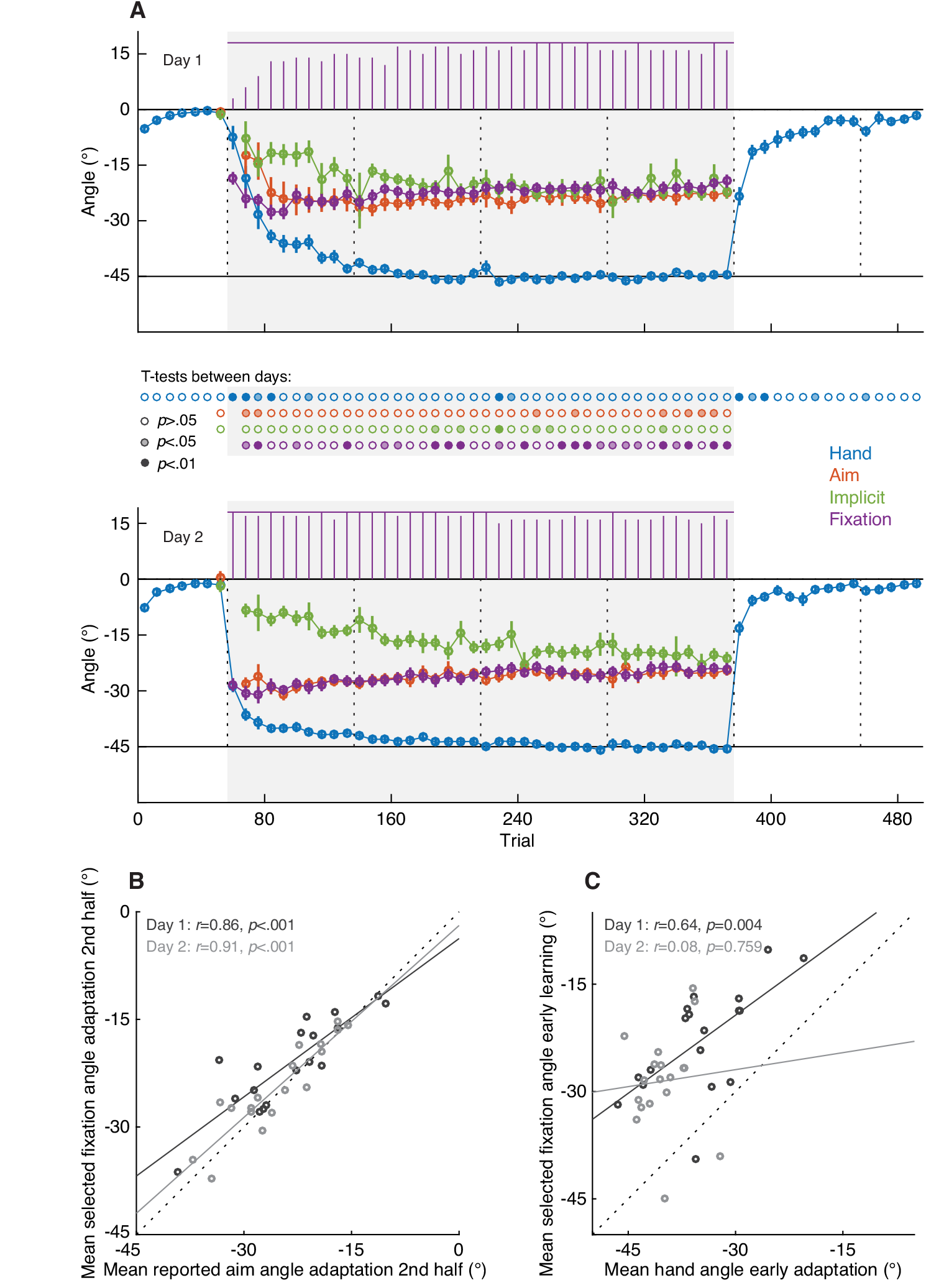
Results Intermittent Report Experiment. **A.** Endpoint hand angles (blue), reported aim angles (orange), implicit angles (green), and selected fixation angles (purple) on day 1 (top) and day 2 (bottom), averaged across aimpoint fixators (n=18) in Experiment 1. Each data point represents the average of a set of eight trials, with error bars showing ±1 SEM across subjects. Purple bars at the top of each graph depict the number of participants contributing to the average selected fixation angle in each trial set. The grey background indicates when the 45° rotation was applied to the cursor feedback. Vertical dotted lines indicate the timing of 30 s breaks during the experiment. The rows of dots in between the top and bottom graphs show the results of uncorrected paired t-tests between each of the data points on day 1 and 2, with the color saturation indicating the significance level. **B**. Relation between the reported aim angle and the selected fixation angle, averaged across the second half of the rotation block of day 1 and day 2. Dashed line indicates the unity line. **C**. Relation between hand angle and selected fixation angle during early adaptation (trial sets 2-10 of the rotation block). *R* and *p* values in B and C show Pearson’s correlation coefficient and its significance value, respectively.

#### Estimating the explicit and implicit component

For Experiment 1 (Intermittent Report Experiment) we estimated the explicit component of visuomotor adaptation using the verbally reported aiming direction (Taylor et al., 2014). We converted the verbally reported landmark number to an angle relative to the target. The reported aiming angles were averaged across sets of eight trials. As such, each value per eight-trial set represents the average of two report trials. Subsequently, implicit adaptation was estimated for each set by subtracting the averaged explicit angle from the averaged hand angle (Taylor et al., 2014). Because, in Experiment 1, we found that the aimpoint fixation angle closely matched the explicit, verbally reported aimpoint angle (see Results), for Experiment 2 we estimated implicit adaptation for each trial set by subtracting the averaged aimpoint fixation angle from the averaged hand angle.

#### Statistical analyses

To assess differences in task performance between day 1 and day 2, we performed paired t-tests on the hand angles, reported aiming angles, implicit angles, and fixation angles averaged across sets of eight trials. To assess differences in adaptation between subgroups of aimpoint and target-only fixators, we performed unpaired t-tests on the hand angles averaged across sets. We computed Pearson’s correlation coefficients to assess, across participants, the relationship between variables.

## Results

The goal of our study was to assess whether gaze behaviour, a natural component of reach planning, can be reliably used to probe both the implementation and state of cognitive strategy use during visuomotor rotation learning and relearning 24 hours later. We predicted that gaze fixations, prior to reaching on each trial, would closely track participants’ verbally reported aiming direction, as assayed on separate trials (Experiment 1). Upon establishing this link, we further predicted that gaze fixations, in the absence of any verbal reporting, would provide a unique means of identifying individuals using cognitive strategies (Experiments 2 and 3). In all three experiments, we predicted that gaze behaviour would be directly related to individuals’ rate of visuomotor adaptation and expression of savings.

## Experiment 1: Intermittent Report

The first experiment contained two separate sessions, separated by 24 hours, of baseline reaches with veridical cursor feedback, adaptation to a 45° visuomotor rotation of the cursor feedback, and washout with veridical feedback. During the rotation block, in 25% of trials, participants were asked to report the number of the landmark they planned to aim their hand to for the cursor to hit the target. We extracted patterns of gaze fixations in the remaining 75% of trials. As shown in Figure 2A and B, typical gaze behaviour involved first shifting gaze from the start position to the visual target at about 200 to 300 ms following target onset. Next, gaze often shifted to a position somewhere in between the visual target and the hand target for two to three fixations. Thereafter, gaze typically remained either in the aimpoint area or shifted back to the visual target before the onset of the reach. Figure 2C shows gaze behaviour over the time course of a single trial during the first rotation block as the probability of fixation in the start point area, visual target area, and the area in between the visual target and the hand target at -45° (‘aim area’). Probabilities were averaged across the subgroup of 18 participants that showed a bimodal pattern of fixation angles (‘aimpoint fixators’; see below).

Figure 3A and B show, for two example participants, the raw endpoint hand angles (in blue), reported aiming angles (in red), and the angles of all fixations during the target preview period of non-reporting trials (in purple). The participant depicted in Figure 3A shows rapid adaptation of the endpoint hand angle from 0° to -45°, with quicker adaptation on the second day compared to the first day (i.e., savings). Further, their verbally reported aiming angle shows a similarly fast change towards -45° in the beginning of the rotation block, and then very slowly drifts back towards about -20° by the end of the rotation block. This participant shows gaze fixations both at the visual target and at an angle in between the visual target and the hand target. The tail of the distribution, denoted by the darker colored purple dots that show the selected fixation angle closest to the hand target at -45° in each trial, very closely mimics the temporal evolution of verbally reported aiming angles during the task. Notably, during the washout block, this participant seems to fixate an additional ‘aimpoint’ location at the diametrically opposite side of the target, as if the rotation were reversed rather than turned off. Many participants exhibited this same behaviour, suggesting that reversion to baseline during washout involves the implementation of a reverse strategy. The right column of Figure 3A shows that the histogram of fixation angles during the rotation block for this participant was well fit by a mixture of two Gaussian curves. When we applied this same approach to the histogram of fixation angles of each participant (see Methods), we found that for 18 out of our 21 participants the histogram was better fit by a mixture of two Gaussian curves than one Gaussian curve, and we thus classified these individuals as ‘aimpoint fixators’. Only two participants showed fixations almost exclusively at the visual target location (i.e., were poorly fit by a mixture of two Gaussian curves), thus classifying these individuals as ‘target-only fixators’. Figure 3B shows the raw hand angles, reported aiming angles, and fixation angles of a target-only fixator. The one, remaining participant switched from a unimodal distribution of fixation angles on day 1 to a bimodal distribution on the second day. To verify our approach of describing the time course of aimpoint fixations by selecting, on each trial, the fixation angle closest to the hand angle, we performed a correlation, across participants, between the mean of the selected fixation angles and the mean of the Gaussian curve in the aimpoint area. This analysis revealed a linear relationship across aimpoint fixators, with a slope close to one on day 1 (slope=1.19, 95% CI=[1.02 1.36], intercept=8.82, 95% CI=[5.06, 12.58]) and day 2 of testing (slope=0.99, 95% CI=[0.84 1.13], intercept=2.13, 95% CI=[-1.62 5.86]).

Figure 4A shows the endpoint hand angles, reported aiming angles, implicit adaptation angles obtained by subtraction of the reported aiming angles from the hand angles, and fixation angles closest to the hand target during the target preview period. All angles are averaged across sets of eight trials and across subjects classified as aimpoint fixators (i.e., 18 out of 21 participants). The time course of gaze fixations closely overlapped with that of the reported aiming angle, confirming our initial hypothesis that gaze fixations would closely track participants’ verbally reported aiming direction. To directly assess the relationship between these two variables, we computed correlation coefficients on the reported aiming angles and the fixation angles on each day, averaged across the trial sets in the second half of the rotation block (i.e., trials 209-368). We observed a strong linear relationship between mean reported aiming angles and mean fixation angles on both days (Figure 4B).

As a descriptive analysis of differences between testing days, we performed paired t-tests between each of the data points on day 1 and 2 (uncorrected for multiple comparisons). Consistent with prior work (e.g., Krakauer et al., 2005; Morehead et al., 2015), we found faster adaptation of the hand angle in the rotation block and washout block of day 2 compared with day 1 (i.e., savings). Notably, faster changes in hand angle following the onset of the rotation on day 2 were accompanied by a larger (i.e., more negative) reported aiming angle, as well as a larger fixation angle, without significant differences in the implicit angle. This suggests that savings were mainly driven by recall of an aiming strategy, possibly facilitated by gaze fixations. [Note that fixation angle differed significantly between days in several bins of the rotation block, however, there was no clear pattern in these differences with the exception that they generally reflected the tendency for aimpoint fixations to be magnified on day 2.] To directly assess the relationship between learning and fixation angle, we computed a correlation, across participants, between the mean hand angle and mean fixation angle during early learning on both days (i.e., trial sets 2-10 of the rotation block). As shown in Figure 4C, this revealed a positive linear relationship on day 1, but not on day 2. We suspect that the lack of a correlation on day 2 might reflect the fact that, due to day 1 learning, the variability across subjects in hand angles was much smaller on day 2 than on day 1.

Taken together, the main results of this Intermittent Report experiment are that (1) the vast majority of participants fixated an internal aimpoint, used to counteract the rotation, prior to executing the reach movement, (2) the magnitude and time course of these ‘aimpoint fixations’ closely overlapped with that of the verbally reported aiming angle, and (3) a greater aimpoint fixation angle during early learning on day 1 was related to greater changes in hand angle.

## Experiment 2: No Report

One plausible explanation of the Experiment 1 findings is that, because we asked participants to verbally report (and thus, presumably fixate) their aimpoint on a minority (25%) of trials, this may have unintentionally biased their gaze patterns on the remaining majority (75%) of trials. To assess whether the nature of the verbal reporting task biased the resulting eye movement patterns, a second group of participants performed the same two sessions of the visuomotor rotation task but without the requirement to report their aiming direction. Here, we found that two subgroups of participants clearly emerged. Eleven out of our 21 participants were now classified as aimpoint fixators based on the fitted Gaussian curves. As in Experiment 1, we again observed a strong linear relationship between the mean of the Gaussian curve in the aimpoint area and the mean selected fixation angle (day 1: r=0.82, p=.002; day 2: r=0.93, p<.001). Notably, the proportion of aimpoint fixators in this experiment was significantly less than in Experiment 1 (Pearson Chi-Square(1)=5.27, p=0.022). In addition, we now found that 8 participants exhibited fixations only around the visual target, and two participants switched from only fixating the target on day 1 to fixating both the target and an aimpoint on day 2 (excluded from the analysis).

Figure 5A shows the endpoint hand angles, fixation angles and the implicit angles estimated by subtracting the fixation angles from the hand angles, averaged across the subgroup of 11 aimpoint fixators, as well as the hand angles averaged across the subgroup of 8 target-only fixators. Whereas the subgroup of aimpoint fixators exhibited adaptation rates that were very similar to those of the aimpoint fixators in Experiment 1 (see *Comparison between Experiments 1 and 2* below), adaptation rates in the subgroup of target-only fixations were considerably slower, with significant between-group differences in hand angles in several sets of trials in the rotation block and early in the washout block. Moreover, whereas the subgroup of aimpoint fixators showed savings, as indicated by significantly faster adaptation on day 2 compared to day 1 in the first trial set of the rotation (t(10)=4.33, p=.002) and washout blocks (t(10)=-3.48, p=.006), the subgroup of target-only fixators failed to show significant savings (first set in rotation block: t(7)= 1.87, p=0.104; first set in washout block: t(7)=-1.76, p=.122). This suggests that learning in the target-only fixators group was largely implicit and did not involve an aiming strategy. To directly assess the relationship between learning and fixation angle, we computed a correlation, across participants, on the mean hand angle and mean fixation angle during early learning (i.e., sets 2-10 of the rotation block). We observed a strong positive correlation on day 1, and a moderate, marginally significant correlation on day 2 (Figure 5B).

**Figure 5.**
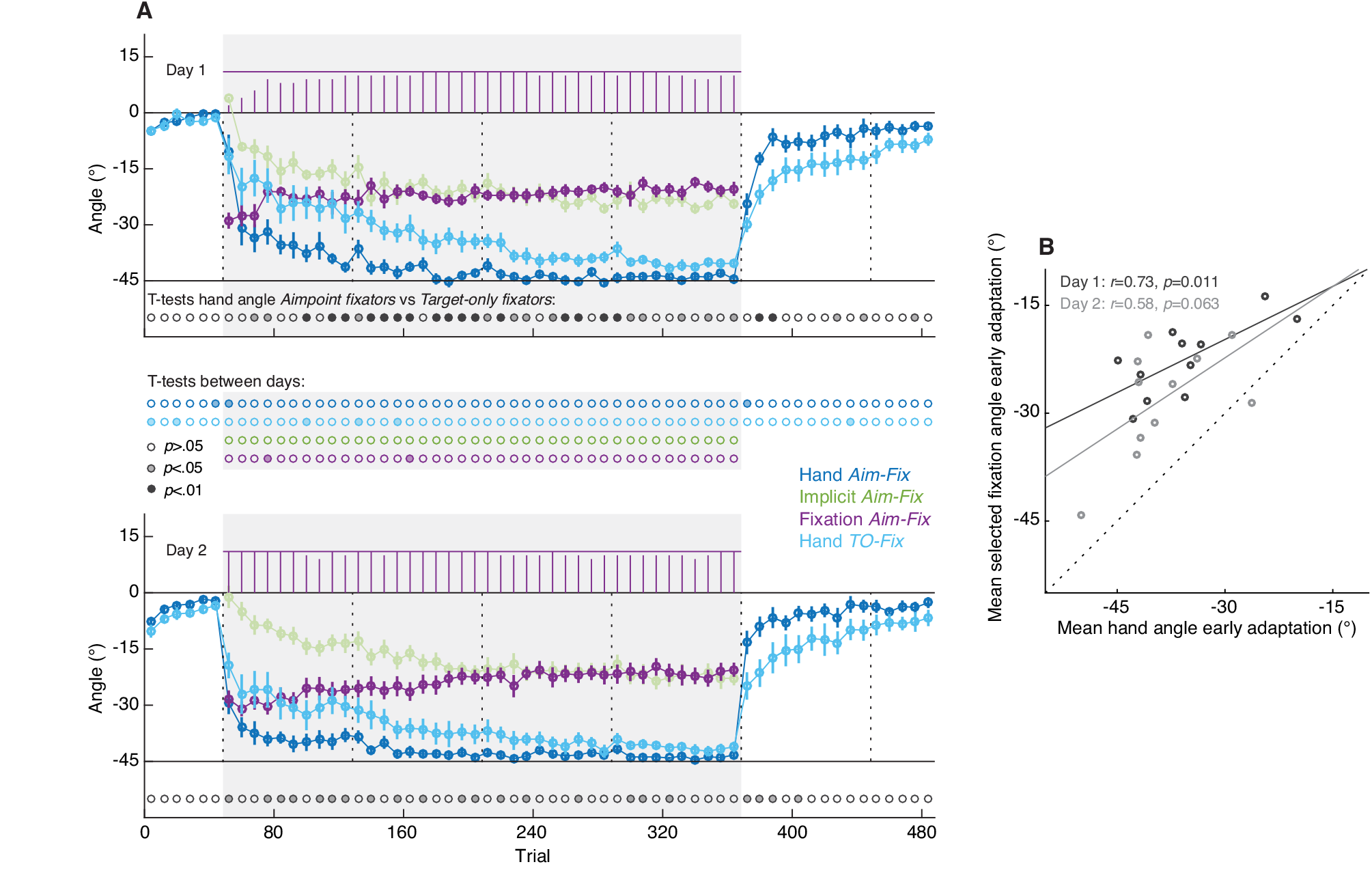
Results No Report Experiment. **A**. Endpoint hand angles, implicit angles (estimated through subtraction of fixation angles from hand angles), and selected fixation angles, averaged across aimpoint fixators (*Aim-Fix,* n=11), as well as endpoint hand angles averaged across target-only fixators *(TO-Fix,* n=8) in Experiment 2. Each data point represents the average of a set of eight trials, with error bars showing ±1 SEM across subjects. Purple bars at the top of each graph show the number of aimpoint fixators contributing to the average selected fixation angle. The grey background indicates when the 45° rotation was applied to the cursor feedback. Vertical dotted lines indicate the timing of 30 s breaks during the experiment. The row of dots at the bottom of each graph shows the result of unpaired t-tests between the aimpoint fixators and the target-only fixators. The rows of dots in between the top and bottom graphs show the results of uncorrected paired t-tests between each of the data points on day 1 and 2, with the color saturation indicating the significance level. **B**. Relation between hand angle and selected fixation angle during early adaptation (trial sets 2-10 of the rotation block). *R* and *p* values show Pearson’s correlation coefficient and its significance value, respectively.

Taken together, the results from this second experiment suggests that (1) the use of verbal reporting measures increases the proportion of participants that implement cognitive strategies, and (2) participants who naturally exhibit aimpoint fixations show fast adaptation and savings whereas those participants who only ever exhibit target fixations show comparably slow adaptation, with no evidence for savings.

## Comparison between Experiments 1 and 2

Across the first two experiments, we found that a significantly larger proportion of participants fixated an aimpoint prior to reaching under a visuomotor rotation when the task involved verbally reporting the aiming direction on a subset of trials. When we compared the hand angle of all subjects in the Intermittent Report experiment (Experiment 1) to the hand angle of all subjects in the No Report experiment (Experiment 2; data not shown), we found significant differences in a large part of the rotation block, especially on the first day of testing. On average, participants in the Intermittent Reporting experiment showed faster adaptation and deadaptation and a greater asymptotic adaptation level than participants in the No Report experiment. However, when we compared the subgroups of aimpoint fixators in both experiments (18 participants in Experiment 1 and 11 participants in Experiment 2), there were no significant differences in hand angle, except in two out of the 55 bins across the entirety of the rotation and washout blocks of each day. These results suggest that the declarative nature of verbal reporting increases the proportion of participants that implement an aiming strategy, resulting in faster learning, but does not affect the magnitude of the explicit component.

## Experiment 3: No Preview

To assess whether a brief (2 s) preview period of the target is necessary for aimpoint fixations to occur, we performed a third experiment. This No Preview experiment was identical to the No Report experiment (Experiment 2), with the exception that, on each trial, participants were instructed to initiate a reach movement upon appearance of the target (i.e., no preview period), and participants performed only a single session of the visuomotor rotation task. Visual inspection of the fixation angles revealed that, despite the lack of a target preview period, five out of twelve participants still showed fixations in the area between the visual target and the hand target during of the rotation block. Figure 6A and B show the raw hand angles, fixation angles, and hand reaction times of two such participants. One participant (Figure 6A) appeared to implement an aiming strategy about half way through the rotation block whereas the other (Figure 6B) implemented an aiming strategy at the start of the rotation block. Unlike in the first two experiments, however, aimpoint fixations generally did not persist throughout the entire rotation block, but were only expressed at what appears to be the start of the implementation of a aiming strategy, as judged from corresponding fast changes in hand angle. As can be seen in the third column of Figure 6A and B, fixating an aimpoint came at the cost of a higher reaction time. The right column of Figure 6A and B show the relation between the selected fixation angle and the hand reaction time for both participants. On average, the aimpoint fixators showed a significant negative relationship between selected fixation angle and reaction time of the hand movement (mean±SEM *r*=-0.34±0.07, one-sample t-test against zero *t*(4)=-4.95 p=.008).

**Figure 6.**
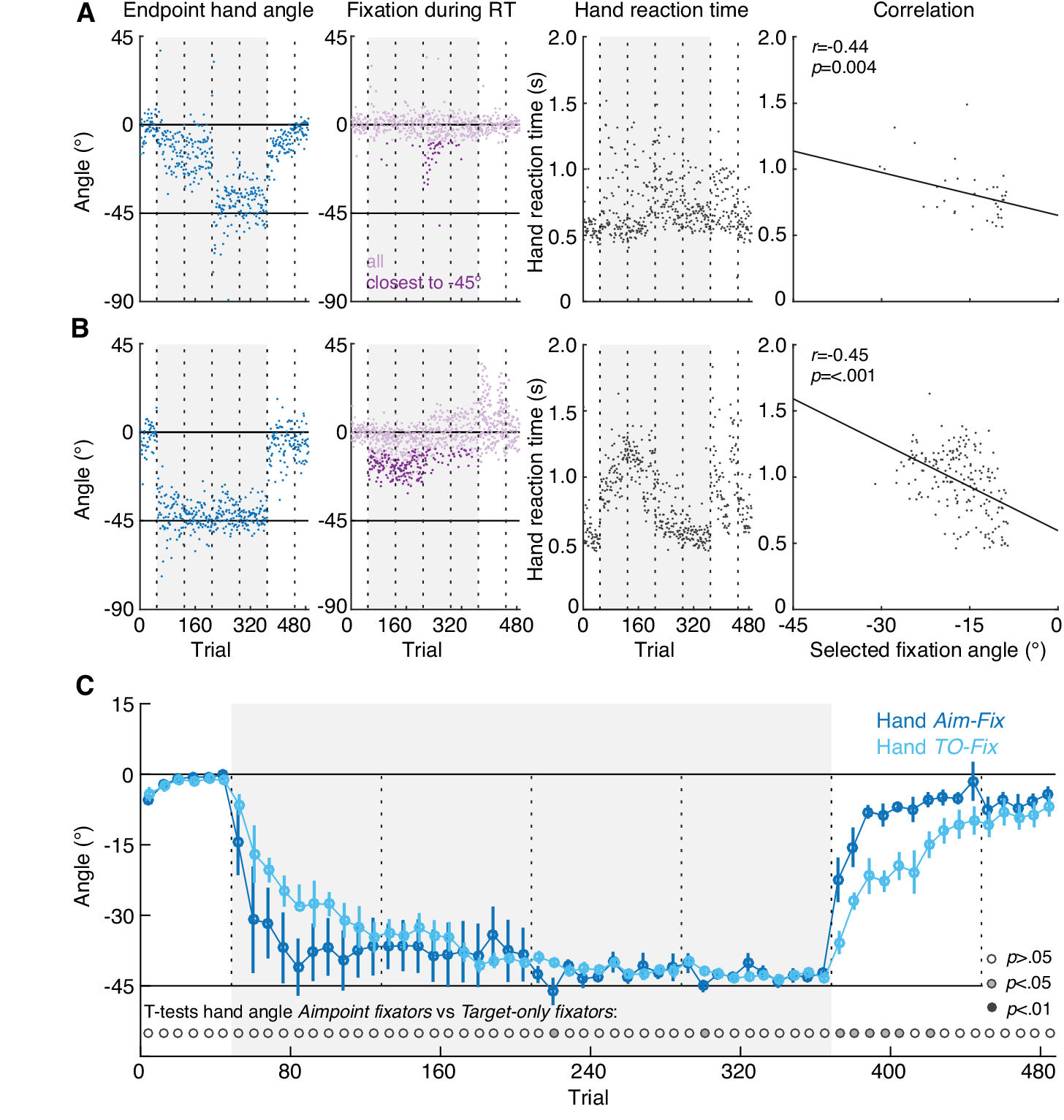
Results No Preview Experiment. **A-B**. Raw endpoint hand angles (blue), fixation angles during the hand reaction time interval (RT; purple), and hand reaction times (black) of two example aimpoint fixators (A and B) in Experiment 3. The grey background indicates when a 45° rotation was applied to the cursor feedback. Vertical dotted lines indicate the timing of 30 s breaks during the experiment. The darker purple dots show, for each trial, the selected fixation angle closest to the hand target. The rightmost column shows the relation between selected fixation angles and hand reaction time. *R* and *p* values show Pearson’s correlation coefficient and its significance value, respectively. **C**. Endpoint hand angles averaged across aimpoint fixators (A*im-Fix,* n=5) and target-only fixators (*TO-Fix,* n=7) in Experiment 3. Each data point represents the average of a set of 8 trials, with error bars showing ±1 SEM across subjects. The row of dots at the bottom of the graph shows the result of unpaired t-tests between the aimpoint fixators and the target-only fixators.

Figure 6C shows the endpoint hand angles averaged across the subgroup of five aimpoint fixators and seven target-only fixators. As in Experiment 2, adaptation and washout were faster for the aimpoint fixators than for the target-only fixators, although this difference was only statistically significant early in the washout block. The lack of significant differences in the rotation block was due to the example participant shown in Figure 6A, who started off as a slower learner, but halfway through the rotation block began implementing an aimpoint fixation strategy, thereby quickly reducing their errors.

To summarize, we found that even without a brief, instructed preview period of the visual target, nearly half the participants still fixated an internal aimpoint location, which, while resulting in faster adaptation, came at the cost of longer reaction times. Notably, in these aimpoint fixators, fixations further away from the visual target (i.e., a greater aiming angle) were associated with longer hand reaction times, consistent with the idea that explicit aiming may involve the mental rotation of a movement endpoint or trajectory (Anguera et al., 2010; Fernandez-Ruiz et al., 2011; McDougle and Taylor, 2016). This experiment reinforces the findings from the first two experiments that trial-to-trial gaze behaviour during adaptation is linked to the implementation of a cognitive strategy.

## Discussion

Here we explored the idea that task-specific gaze fixations over the time course of sensorimotor adaptation and readaptation 24 hours later can provide a covert means of identifying individuals who use explicit strategies during learning, as well as how the contribution of the explicit component to learning evolves over time. We show, across three experiments, that gaze behaviour during visuomotor rotation learning parcellates the explicit and implicit components to learning, is linked to individual differences in learning rates, and can predict the expression of savings.

Previous research has examined natural gaze behaviour during adaptation to a visuomotor rotation in the presence of visual landmarks. This study showed that without online cursor feedback, gaze behaviour shifted from being slightly more often directed to the vicinity of the visual target, around the time of hand movement onset, during early learning, to being more often directed to the vicinity of the hand target during late learning (Rand and Rentsch, 2016). Although this finding suggests a possible relation between gaze behaviour and reaiming strategies, it is inconsistent with the well-characterized fast increase and then gradual reduction of the explicit component during adaptation (e.g., current study; Taylor et al., 2014; McDougle et al., 2015). We suspect that some of the discrepancy between this prior work and our current findings reflects the fact that this prior work focused on (1) gaze locations during single time intervals, which as we have shown is probabilistic rather than circumscribed (see Figure 2C), and, (2) an analysis of group data. Our current results are able to directly link gaze fixations, prior to hand movement, to the explicit component of learning and savings in a trial-by-trial fashion and individual participant basis.

### Gaze behaviour as a substitute to verbal reporting

The large participant groups tested in the current study allowed us to divide individuals into two main subgroups: (1) a group that only fixated the visual target (i.e., target-only fixators) and, (2) a group that fixated both the visual target and a separate aimpoint (i.e., aimpoint fixators). Our results indicate that group membership is affected by verbal reporting, such that the requirement to declare aiming direction on a subset of trials increases the number of aimpoint fixators rather than the magnitude of the explicit component, as previously assumed (Taylor et al., 2014; Leow et al., 2017). We found that target-only fixators adapted more gradually, and did not exhibit savings, indicative of implicit processes governing their learning and relearning of the visuomotor rotation (Morehead et al., 2015). By contrast, aimpoint fixators exhibited fast adaptation and savings, indicating the use of explicit strategies.

We further noticed that several participants were quite rigid in their verbal reporting; that is, they consistently tended to report, across trials, a fixed number of landmarks counterclockwise to the visual target as their aimpoint. The declarative nature and rigidness of reporting suggests an advantage to using gaze to assess the contribution of explicit processes to learning. First, the lack of aimpoint fixations may identify participants who, in the absence of being prompted by verbal reporting, would not spontaneously implement an aiming strategy. Second, in participants who do implement such strategies, gaze can provide a covert, yet sensitive measure of the magnitude of the explicit component.

We recognize, however, that there may also be some shortcomings in using gaze fixations to assess the explicit component. First, the absence of aimpoint fixations does not preclude the possibility that explicit strategies are still being implemented. However, the gradual nature of learning and absence of savings in non-aimpoint fixators suggests that their learning is largely implicit (Morehead et al., 2015). Second, adding landmarks to the visual scene and providing a brief target preview are two simple, but likely essential, modifications needed to robustly elicit aimpoint fixations in participants that naturally implement a cognitive strategy. Nevertheless, given that the primary method used for dissociating the explicit and implicit components to learning involves declarative reporting (Taylor et al., 2014), which itself enhances the probability that cognitive strategies are implemented and which also necessitates the use of landmarks and increases reaction times, we believe that the use of gaze behaviour has inherent advantages.

### The role of aimpoint fixations during visuomotor learning

During a trial, aimpoint fixators typically shifted their gaze from the start position to the visual target shortly after its appearance, and then shifted their gaze (in one or a series of saccades) to the aimpoint. Presumably, aimpoint fixations assist participants in performing a mental rotation of the motor goal location or movement direction (McDougle and Taylor, 2016). This suggestion is supported by our observation that in the No Preview experiment, aimpoint fixators’ hand reaction times were correlated with the magnitude of their fixation angles. This relationship between rotation magnitude and reaction time bears strong similarity to previous observations in studies of visually guided reaching and object rotation (Shepard and Metzler, 1971; Pellizzer and Georgopoulos, 1993).

When reaching under normal visual feedback conditions, fixating the target optimizes the use of peripheral visual feedback in automatically correcting for errors in the reach trajectory (e.g., Carlton, 1981; Paillard, 1996; Land et al., 1999; Saunders and Knill, 2003; de Brouwer et al., 2017). Although in the current study participants were instructed to make ballistic, uncorrected reaching movements, it is unlikely that participants fully ignored peripheral visual information of the cursor, which provides an important reason for fixating the target. Surprisingly, however, aimpoint fixators were more likely to keep their gaze at the aimpoint location than shift it back to the visual target location during the actual execution of the reach (52% vs. 40% probability, respectively). One explanation for this behaviour is that fixating the hand target could improve reach accuracy through the use of extraretinal signals; that is, proprioceptive signals or an efference copy of oculomotor commands (e.g., Prablanc et al., 1986). It is important to recognize, however, that participants did not actually direct their gaze to the hand target (which would, in principle, provide the most spatially accurate extraretinal information to hit the target), but rather a strategic location to counteract the rotation. One intriguing possibility, which may explain why gaze often remained at this location, is that the trial-by-trial state of the implicit component during learning is directly built into the transformation from gaze proprioceptive coordinates to the hand movement. Although previous work has examined reference frame transformations from gaze-centred to hand-centred coordinates (Crawford et al., 2004; Buneo and Andersen, 2006), it has not directly explored how this mapping might be affected by implicit learning.

### Gaze behaviour during washout

Strikingly, we observed that almost all participants who fixated an aimpoint during the rotation block also appeared to fixate an aimpoint, in the opposite direction, in the deadaptation (washout) blocks on both days. That is, even though veridical visual feedback was restored during washout, the distribution of gaze angles appeared to be bimodal with a second peak in between the visual target and +45°, as if the rotation were reversed rather than extinguished. This indicates that deadaptation itself also involves an explicit component and not just the gradual reduction of the implicit component, for which it is commonly used (e.g., Krakauer et al., 2005). This finding is consistent with recent work showing that deadapting to an instantaneously removed rotation, A, results in savings when subsequently experiencing rotation -A (Herzfeld et al., 2014). The idea that deadaptation involves an explicit component appears to contradict recent findings from Morehead and colleagues (2015) who asked participants to verbally report their aiming direction during deadaptation and found that participants aimed towards the target rather than an opposite aimpoint. This discrepancy might be due to differences in the magnitude of the implicit component at the time the rotation was removed. Namely, when the implicit component at the end of the rotation block is small, as in the Morehead study (~10°), extinguishing the rotation will produce only a small error between the target and the cursor position, which is less likely to drive an aiming strategy (Bond and Taylor, 2015). Notably, for many participants in our study, gaze remained at an ‘opposite’ aimpoint throughout the full 120 trials of deadaptation, suggesting that the implicit component was not, in fact, washed out (as explicit aiming was being used to counteract it). Further research is needed to carefully unravel the complete time course of the explicit and implicit components to deadaptation.

### Brain mechanisms linking gaze and explicit processes

Whereas there is extensive evidence that implicit, error-based sensorimotor adaptation is reliant on cerebellar mechanisms (Smith and Shadmehr, 2005; Morton and Bastian, 2006; Tseng et al., 2007), the neural systems associated with the explicit component of learning remain largely unknown. The verbal reporting task employed by Taylor et al. (2014) showed that the use of explicit strategies in VMR learning can be declarative. Although Experiment 2 did not involve verbal reporting, we suspect that aimpoint fixators, if queried, would similarly acknowledge use of such strategies. Perhaps not surprisingly, evidence from neuroimaging, aging, and lesion studies, has implicated prefrontal cortex in explicit strategies (Taylor and Ivry, 2014). Several studies have implicated dorsolateral prefrontal cortex, in particular, in contributing to sensorimotor adaptation and savings (Shadmehr and Holcomb, 1997; Della-Maggiore, 2004; Floyer-Lea and Matthews, 2004), likely through its known role in working memory processes (Curtis and D’Esposito, 2003; Seidler et al., 2012) and mental rotation (Cohen et al., 1996). With respect to the current results, we expect the frontal eye fields, located in prefrontal cortex and a key hub in the oculomotor network associated with target selection (Thompson and Bichot, 2005), to be involved in the selection of aimpoints as saccade targets. The role of declarative processes in strategic re-aiming further suggests that regions in the medial temporal lobe (MTL) might also be partly responsible for the reported oculomotor behaviour. MTL regions appear integral to guiding gaze to strategic locations in a visual scene (Meister and Buffalo, 2016) and the neuroanatomical connectivity of the MTL makes it well poised to interface with oculomotor regions in prefrontal cortex (Shen et al., 2016).

